# CAR T cell entry into tumor islets is a two-step process dependent on IFNγ and ICAM-1

**DOI:** 10.1101/2020.05.27.119198

**Authors:** Chahrazade Kantari-Mimoun, Sarah Barrin, Sandrine Haghiri, Claire Gervais, Joerg Mittelstaet, Nadine Mockel-Tenbrinck, Ali Kinkhabwala, Diane Damotte, Audrey Lupo, Mathilde Sibony, Marco Alifano, Lene Vimeux, Elisabetta Dondi, Nadège Bercovici, Alain Trautmann, Andrew Kaiser, Emmanuel Donnadieu

## Abstract

Adoptive transfer of T cells expressing chimeric antigen receptors (CARs) has shown remarkable clinical efficacy against advanced B cell malignancies but not yet against solid tumors. Here, we used fluorescent imaging microscopy and *ex vivo* assays to compare the early functional responses (migration, Ca^2+^ and cytotoxicity) of CD20 and EGFR CAR T cells upon contact with malignant B and carcinoma cells. Our results indicate that CD20 CAR T cells rapidly form productive ICAM-1-dependent conjugates with their targets. By comparison, EGFR CAR T cells only interact at first with a subset of carcinoma cells located at the periphery of tumor islets. After this initial peripheral activation, EGFR CAR T cells progressively infiltrate the center of tumor cell regions. The analysis of this two-step entry process shows that activated CAR T cells trigger the upregulation of ICAM-1 on tumor cells in an IFNγ-dependent pathway. The blockade of ICAM-1/LFA-1 interactions prevents CAR T cell recruitment into tumor islets. The requirement for IFNγ and ICAM-1 to enable CAR T cell entry into tumor islets is of significance for improving CAR T strategy in solid tumors.

## INTRODUCTION

Adoptive immunotherapy with gene-engineered chimeric antigen receptor (CAR) T-cells has demonstrated its striking efficacy in several B cell malignancies (1). Two CD19 CAR T cells, Kymriah™ and Yescarta™ have obtained in 2017 the FDA approval. However, the field of CAR T cells is currently facing two major challenges (2). First, although CAR T cells can be extremely effective in killing malignant cells, they also can cause dangerous side effects including off-tumor toxicity and cytokine release syndrome (3). The second major challenge is that the CAR T cell strategy has not yet produced favorable clinical responses in solid malignancies (4). This could be due at least in part to the existence of various physical and environmental barriers in solid tumors, which are less present in hematological cancers. In solid tumors, lymphocytes have to home and go through a dense stroma that surrounds cancer cells. Once within the tumor, T cells must expand, persist, and mediate cytotoxicity in a hostile environment largely composed of immunosuppressive cells and molecules. Using an experimental system of human tumor slices combined with dynamic imaging microscopy, we have previously demonstrated that a dense extracellular matrix (ECM) (5, 6) and tumor-associated macrophages (TAM) (7) hinder anti-tumor functions of T cells in progressing tumors, by reducing their migration and interaction with tumor cells. After crossing these stromal obstacles, CD8 T cells need to infiltrate tumor islets to produce their cytotoxic activity. Up to now, little is known about the elements that promote CAR T cell interaction with their targets. Adhesion molecules among which two integrins, namely lymphocyte function-associated antigen-1 (LFA-1) and CD103 (αE/β7), play a critical role in T - tumor cell interaction (8, 9). LFA-1 (also known as CD11a/CD18 and αLβ2) is an integrin expressed by T cells and other hematopoietic cells. This integrin binds to intercellular adhesion molecule 1 (ICAM-1 or CD54) expressed by antigen-presenting cells as well as by some tumor cells. When T cells are stimulated by a variety of external stimuli, including antigens and chemokines, the affinity and clustering of LFA-1 increases in an inside-out signaling process that promotes the binding of this integrin to ICAM-1 (10). Whether similar molecules and mechanisms control the anti-tumoral action of CAR T cells remains to be established.

Here, we used fluorescent imaging microscopy and *ex vivo* assays to compare the activation and infiltration of CD20 and EGFR CAR T cells upon contact with malignant B and carcinoma cells. We describe for the first time a two-step entry of EGFR CAR T cells in human tumor islets of carcinomas. This entry, which is dependent on both IFNγ and ICAM-1 controls the infiltration of EGFR CAR T cells in tumor islets, and the subsequent killing of tumor cells.

## RESULTS

### CD20 CAR T cells form stable conjugates with B malignant cells and rapidly increase their intracellular Ca^2+^

First, we used fluorescent imaging microscopy to visualize the activity of CD20 CAR T cells when contacting CD20 positive tumor cells. CD20 CAR T cells, loaded with the fluorescent dye fura-2, were added onto a monolayer of the B lymphoma cell line Raji. The intracellular Ca^2+^ rise in T cells following CAR engagement is one the earliest signs of T lymphocyte activation.

As shown in **Figure 1A and Movie S1**, the formation of stable conjugates by CD20 CAR T cells with Raji B cells is rapidly followed by large and sustained Ca^2+^ increases. By comparison, untransduced T cells transiently interact with several Raji cells without increasing their Ca^2+^. On average, 80% of CD20 CAR T cells exhibit Ca^2+^ responses induced by Raji B cells (**Figure 1A, right panel**). The few cells that did not respond are likely untransduced ones since 85% of T cells were positive for the CAR.

**Figure 1:**
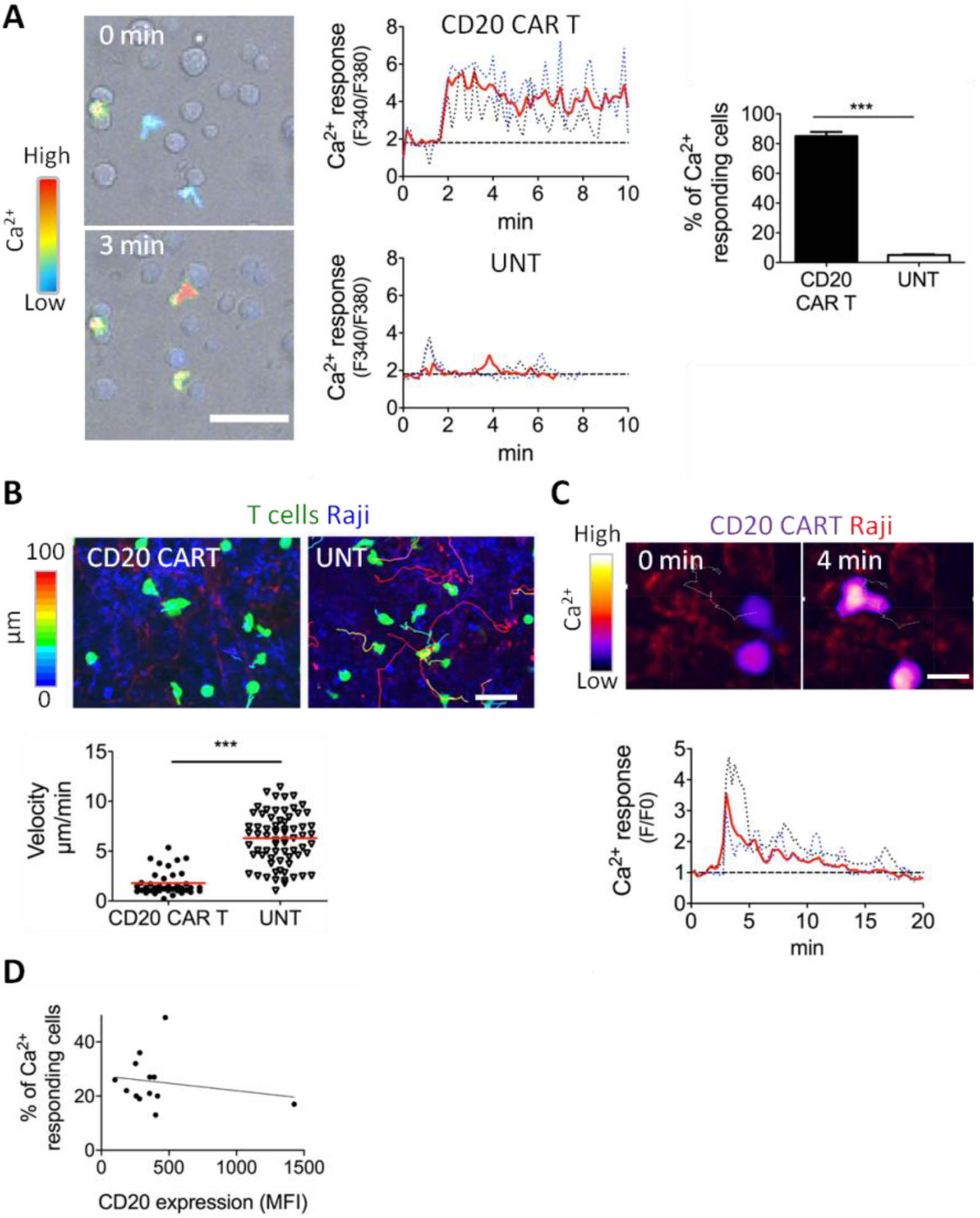
CD20 CAR T cells form rapid productive conjugates with malignant B cells. **(A)** Ca^2+^ response of CD20 CAR T cells contacting Raji B cells. Raji B cells were placed on glass coverslips to adhere for 30 min at 37°C and then washed. CD20 CAR T cells loaded with fura 2-AM were added before image recording. **(A, left)** Snapshots of a time lapse showing Ca^2+^ increases in CD20 CAR T cells after their contacts with Raji B cells. The CAR T cell Ca^2+^ level is coded in color, ranging from blue to red. Scale bar: 50 μm. See also **Movie S1**. **(A, middle)** Ca^2+^ levels of two single CD20 CAR and untransduced T cells (dotted blue lines) plotted against time. Red thick lines show average Ca^2+^ responses (15-20 cells.) Basal Ca^2+^ levels are indicated by black dotted lines. **(A, right)** Percentage of Ca^2+^ responding CD20 CAR T cells among cells in contact with Raji B cells. Results are shown as mean ± SEM; n = 100-150 cells/condition from 5 independent experiments; Student test: ***P < 0.001. **(B-C)** Migration and Ca^2+^ responses of CD20 CAR and untransduced T cells in vibratome sections of viable Raji tumors. (**B, upper)** Snapshots of a time lapse with trajectories of individual CD20 CAR T cells (green) in slices stained with anti-CD19 (blue) Abs for Raji tumor cells. Tracks are color-coded according to CAR T cell displacement length. Scale bar: 30 μm. **(B, lower)** Average speed of CD20 CAR T and untransduced T cells in Raji Tumor slices. Results are shown as mean ± SEM; n = 30-40 cells/condition from 3 independent experiments; Student test: ***P < 0.001. **(C, upper)** Snapshots of a time lapse showing Ca^2+^ response of fluo-4 loaded CD20 CART in vibratome sections of viable Raji tumors at 0 and 4 min. Scale bar: 20 μm. See also **Movie S2. (C, lower)** Ca^2+^ levels of two single CD20 CAR T cells (dotted blue lines) plotted against time. Red thick lines show the average Ca^2+^ response (15-20 cells.) Basal Ca^2+^ levels are indicated by black dotted lines. **(D)** Percentage of Ca^2+^ responding CD20 CAR T cells among cells in contact with B cells from CLL patients (n=13 patients) in relation to CD20 expression at the surface of malignant B cells, determined by flow cytometry.

Next, we monitored the motile behavior and Ca^2+^ responses of CD20 CAR T cells in Raji tumor tissues (**Figure 1B and C**). For these experiments, Raji cells were implanted subcutaneously into immune-deficient mice (NSG). Three weeks later, tumors were vibratome-sliced and CD20 CAR transduced or untransduced T cells were plated onto fresh slices. Whereas untransduced T cells migrated very actively within tumor slices in an apparent random manner without stopping, CD20 CAR T cells were strikingly static, even though not totally arrested (**Figure 1B**). During a 20 min recording, the mean velocity of non-transduced T cells was around 6 μm/min, whereas CD20 CAR T cells exhibited an average velocity of 1.5 μm/min. Unlike untransduced lymphocytes, CD20 CAR T cells exhibited a rapid increase in their intracellular Ca^2+^ levels upon infiltration in Raji tumor slices. With CD20 CAR T cells, a 3.5-fold increase in the intracellular Ca^2+^ level was observed on average upon contact with Raji B cells (**Figure 1C and Movie S2**).

We then monitored Ca^2+^ responses of CD20 CAR T cells during their interaction with malignant B cells from 13 chronic lymphocytic leukemia (CLL) patients. Our results indicate that the percentage of Ca^2+^ responding CAR T cells ranges from 15 to 50%, depending on the patient (**Figure 1D**). This variability was not related to CD20 expression, since there was no correlation between the proportion of activated CD20 CAR T and target antigen density, as assessed by flow cytometry (**Figure 1D**). This result suggests that other surface proteins expressed by tumor cells dictate CD20 CAR T cell responsiveness.

### EGFR CAR T cell responsiveness is dependent on the spatial orientation of carcinoma cells

Imaging experiments were then performed with EGFR CAR T cells and the human tumor pancreatic cell line BxPC3, which express EGFR. Of note, BxPC3 cells did not undergo the epithelial-mesenchymal transition and therefore still express epithelial markers such as EpCAM. In culture, these cells grow in clusters with conspicuous peripheral and central areas (**Figure 2A**).

**Figure 2:**
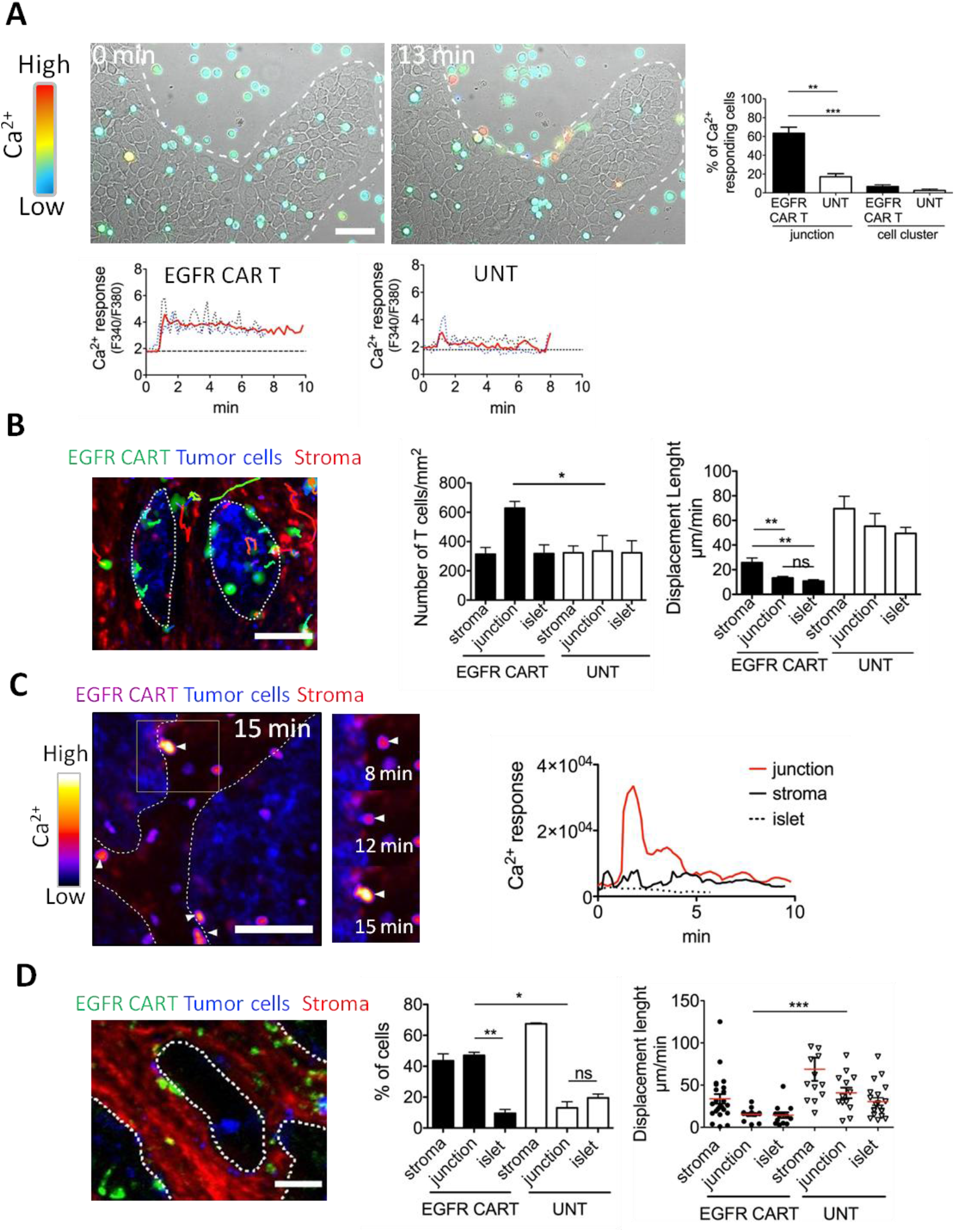
EGFR CAR T cell responsiveness is dependent on the spatial orientation of carcinoma cells. **(A)** Ca^2+^ response of EGFR CAR T cells contacting BxPC3 tumor cells. BxPC3 cells were placed on glass coverslips the day before the experiment. EGFR CAR T cells loaded with fura 2-AM were added before image recording. **(A, upper panel)** Snapshots of a time lapse showing Ca^2+^ increases in EGFR CAR T cells after their contacts with BxPC3 tumor cells. The CAR T cell Ca^2+^ level is coded in color, ranging from blue to red. See also **Movie S3**. **(A, lower panels)** Ca^2+^ levels of two single EGFR CAR and untransduced T cells (dotted blue lines) plotted against time. Red thick lines show average Ca^2+^ responses (15-20 cells.) Basal Ca^2+^ levels are indicated by black dotted lines. **(A, right)** Proportion of Ca^2+^ responding EGFR CAR T and untransduced cells at the periphery or within BxPC3 tumor cell clusters, represented as mean ± SEM; n = 75-100 cells/condition from 4 independent experiments; Student test: **P < 0.01 and ***P < 0.001. (**B)** Concentration and migration of EGFR CAR T cells in BxPC3 tumor slices. **(B, left)** Snapshots of a time lapse with trajectories of EGFR CAR T cells (green) in slices stained with anti-EpCAM (blue) for tumor cells and with anti-Fibronetin (red) for stroma. Tracks are color-coded according to CAR T cell displacement length. White dotted lines delineate tumor islets. See also **Movie S5**. (**B, middle**) Concentration of EGFR CAR and untransduced T cells in the stroma, tumor-stroma junctions and BxPC3 tumor cell regions represented as mean ± SEM; n = 3 independent experiments; Student test: *P < 0.05. (**B, right**) Displacement of EGFR CAR and untransduced T cells in the stroma, tumor-stroma junctions and BxPC3 tumor cell regions represented as mean ± SEM; n = 3 independent experiments; Student test: *P < 0.05. **(C)** Ca^2+^ responses of EGFR CAR T cells in BxPC3 slices. **(C, left)** Snapshots of a time-lapse showing Ca^2+^ response of fluo-4-loaded EGFR CAR T cells in vibratome sections of BxPC3 tumors at 15 min. The boxed areas represented higher magnifications and show snapshots at various time intervals for the zame zone. See also **Movie S6**. (**C, right**) Ca^2+^ levels of one typical EGFR CAR T cell in the stroma, tumor-stroma junction and BxPC3 tumor cell region plotted against time. (**D)** Concentration and migration of EGFR CAR T cells in human lung tumor slices. **(D, left)** Representative images of EGFR CAR T cell distribution in a human lung tumor slice stained for EpCAM (tumor cells) and fibronectin (stroma). White dotted lines delineate tumor islets. See also **Movie S7** (**D, middle**) Proportion of EGFR CAR and untransduced T cells in the stroma, tumor-stroma junctions and tumor islets of human lung tumors, represented as mean ± SEM; n = 3 independent experiments; Student test: *P < 0.05 and **P < 0.01. (**D, right**) Displacement of EGFR CAR and untransduced T cells in the stroma, tumor-stroma junctions and tumor islets of human lung tumors represented as mean ± SEM; n = 3 independent experiments; Student test: ***P < 0.001. Scale bars: 50 μm

We observed that EGFR CAR T cells were not very prone at forming productive conjugates with pancreatic tumor cells (**Figure 2A and S1A)**. Strikingly, the few CAR T cells showing Ca^2+^ increases were those contacting tumor cells located at the periphery of the clusters (**Figure 2A right panel and Movie S3)**. In contrast, few CAR T cells were activated by an interaction with tumor cells in the center of the islets. This differential spatial response was observed with T cells expressing either high (Cetuximab) or low (Nimotuzumab) affinity CARs for EGFR. The only difference between the two CARs was that Ca^2+^ spikes were more oscillatory with the low affinity one (**Figure S1B**). Such a peripheral CAR T cell activation was also noted with the cholangiocarcinoma cell line EGI-1, which shows a spatial organization similar to that of BxPC3 (**Figure S1C)**. The propensity to preferentially activate CAR T cells at the islets outskirts was independent of the target antigen since similar results were obtained with BxPC3 transduced with CD20 and CAR T cells redirected towards CD20 (**Figure S1D and Movie S4**).

We next assessed if this peripheral activation of CAR T cells was observed on fresh slices from BxPC3 tumors isolated from NSG mice. Indeed, EGFR CAR T cells were enriched at the tumor - stroma border, as compared to non-transduced T cells that exhibited a homogenous distribution in all compartments (**Figure 2B and Movie S5**). Moreover, EGFR CAR T cells were relatively static at this border (**Figure 2B right panel**) and made frequent Ca^2+^ responses (**Figure 2C**) in sharp contrast with T cells located in the stroma and in tumor islets (**Movie S6**). On average, 44% of EGFR CAR T cells in contact with tumor cells at the border increased their Ca^2+^ as compared to 22% when lymphocytes were in the center of tumor islets.

Next, we turned to fresh human tumors, to make sure that these findings were not biased by the use of tumor cell lines. The same pattern was evidenced when EGFR CAR T cells were added onto fresh human lung tumor slices, with a structure representative of a carcinoma composed of compact tumor islets surrounded by a fibrous stroma (**Figure 2D left panel**). In terms of CAR T cell distribution, lymphocytes were enriched in the stroma and at the edge of the islets in contact with peripheral tumor cells (**Figure 2D middle panel**). Moreover, CAR T cells in tumor islet periphery were relatively static, compared to untransduced lymphocytes in the same regions (**Figure 2D right panel and Movie S7**). This peculiar enrichment and arrest of EGFR CAR T at the tumor - stroma junction was also confirmed in slices made from human renal cell carcinomas (**Figure S1E and Movie S8**). Although the architecture of renal tumors was slightly different from that of lung tumors similar results were obtained in terms of CAR T cell distribution and displacement length. Next, we performed immunostaining experiments using the MACSima approach to study the distribution of many proteins on a single BxPC3 tumor cryosection. We found that unlike EGFR and EpCAM, which present homogenous distributions, some membrane proteins including CD104, the β4 integrin, exhibit a clear enrichment at the periphery of tumor islets of BxPC3 tumors (**Figure S2**).

Overall, these results reveal that EGFR CAR T cells preferentially form productive conjugates with a minority of carcinoma cells localized at the periphery of the clusters. They also suggest that proteins enriched at the periphery of tumor islets modulate the capacity of the target antigens to be detected by CAR T cells.

### ICAM-1 expression by malignant B cells is instrumental for early CAR T cell activation

The low T cell stimulatory capacity of EGFR-positive carcinoma cells distributed within tumor islets as compared to Raji B cells prompted us to search for surface molecules differentially expressed between both tumor cell types. Our data indicate that ICAM-1 was strongly expressed by the Raji B lymphoma cell line (**Figure 3A**). In sharp contrast, ICAM-1 was hardly detected in tumor islets formed by BxPC3 pancreatic and EG-I1 cholangiocarcinoma tumor cell lines, even at their periphery. To investigate the implication of this adhesion molecule, we measured the Ca^2+^ responses of CD20 CAR T cells induced by malignant B cells from CLL patients expressing various levels of ICAM-1. A clear positive correlation was observed between the percentage of activated CAR T cells and ICAM-1 density (**Figure 3B**). Moreover, blocking anti-ICAM-1 or anti-LFA-1 antibodies markedly decreased the percentage of Ca^2+^ responding CD20 CAR T cells in contact with Raji B cells (**Figure S3**).

**Figure 3:**
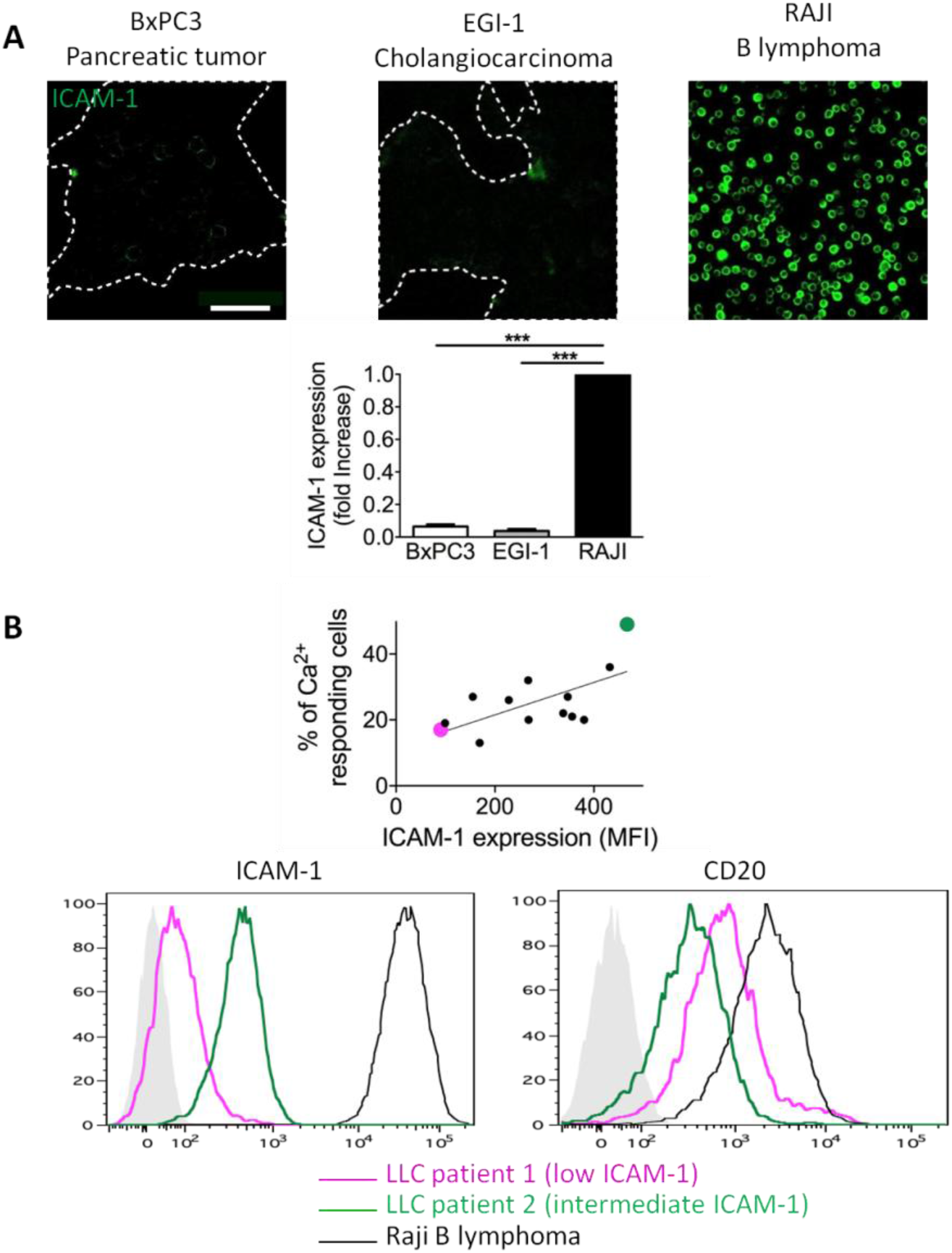
ICAM-1 expression by malignant B cells is instrumental for early CD20 CAR T cell activation. **(A, upper panel)** ICAM-1 expression on pancreatic BxPC3 tumor cells, cholangiocarcinoma EGI-1 cells and Raji B lymphoma cells. Scale bar: 100 μm **(A, lower panel)** Quantification of ICAM-1 expression as determined by image analysis, represented as fold increase. Mean ± SEM; n = 3 independent experiments; Student test: ***P < 0.001. **(B, upper panel)** Percentage of Ca^2+^ responding cells as the function of the ICAM-1 expression at the surface of B cells from malignant CLL (n=13 patients). Pearson coefficient: 0.64. **(C, lower panel)** FACS plot of ICAM-1 and CD20 expression on B cells from 2 CLL patients, also shown in upper panel, and on Raji B lymphoma cells.

These results demonstrate that ICAM-1 expression by tumor cells plays a crucial role in the responsiveness of CAR T cells to their targets. They also suggest that a defect in the expression of ICAM-1 by solid tumors may contribute to explain why EGFR CAR T cells are relatively inefficient in forming productive conjugates with carcinoma cells.

### Activated EGFR CAR T cells mediate ICAM-1 expression by carcinoma cells

ICAM-1 expression is known to be controlled by various inflammatory cytokines, including TNFα and IFNγ, produced by activated T cells (11). We thus wondered if EGFR CAR T cells could induce the upregulation of ICAM-1 on BxPC3 cells. To this end, we performed co-culture experiments between EGFR CAR T cells and BxPC3 carcinoma cells for 4 h at an Effector: Target ratio of 5:1. As shown in **Figure 4A**, ICAM-1 expression on target cells was markedly increased by EGFR CAR T cells but not by untransduced T lymphocytes. Recombinant IFNγ which was used as a positive control partially mimicked the effect of EGFR CAR T cells on ICAM-1 upregulation. Similar results were obtained with the cholangiocarcinoma cells EGI-1 co-cultured with EGFR CART cells for 4 h (data not shown). Conversely, VCAM-1 expression on the surface of BxPC3 tumor cells was not increased by EGFR CAR T cells (**Figure S4A**). In addition, EGFR CAR T cells but not non-transduced T cells were able to induce the phosphorylation of STAT1, a key molecule for IFNγ signalling, in BxPC3 tumor cells (**Figure 4B**). Phopho-STAT1 was detected as early as 1 hour after adding EGFR CAR T cells. CAR T cells were as potent as recombinant IFNγ to induce phospho-STAT1. This result highlights the role of IFNγ produced by activated EGFR CAR T cells in this process. Indeed, we could detect the presence of IFNγ and TNFα in the supernatant of CAR T cell-tumor cell cocultures (**Figure 4C and S4B**). Our results showing that activated CAR T cells drive the expression of ICAM-1 in an IFNγ-dependent fashion led us to investigate the expression of ICAM-1 in human lung and renal carcinomas, in relation with resident T cells. As predicted, in tumors enriched in CD3 T cells, tumor cells strongly express ICAM-1 (**Figure 4D**). In addition, in TCGA databases, a positive correlation may be observed between activated CD8 T cells and ICAM-1 transcripts in human lung squamous cell carcinomas (**Figure 4E and S4C**) and pancreatic adenocarcinomas (**Figure S4D**).

**Figure 4:**
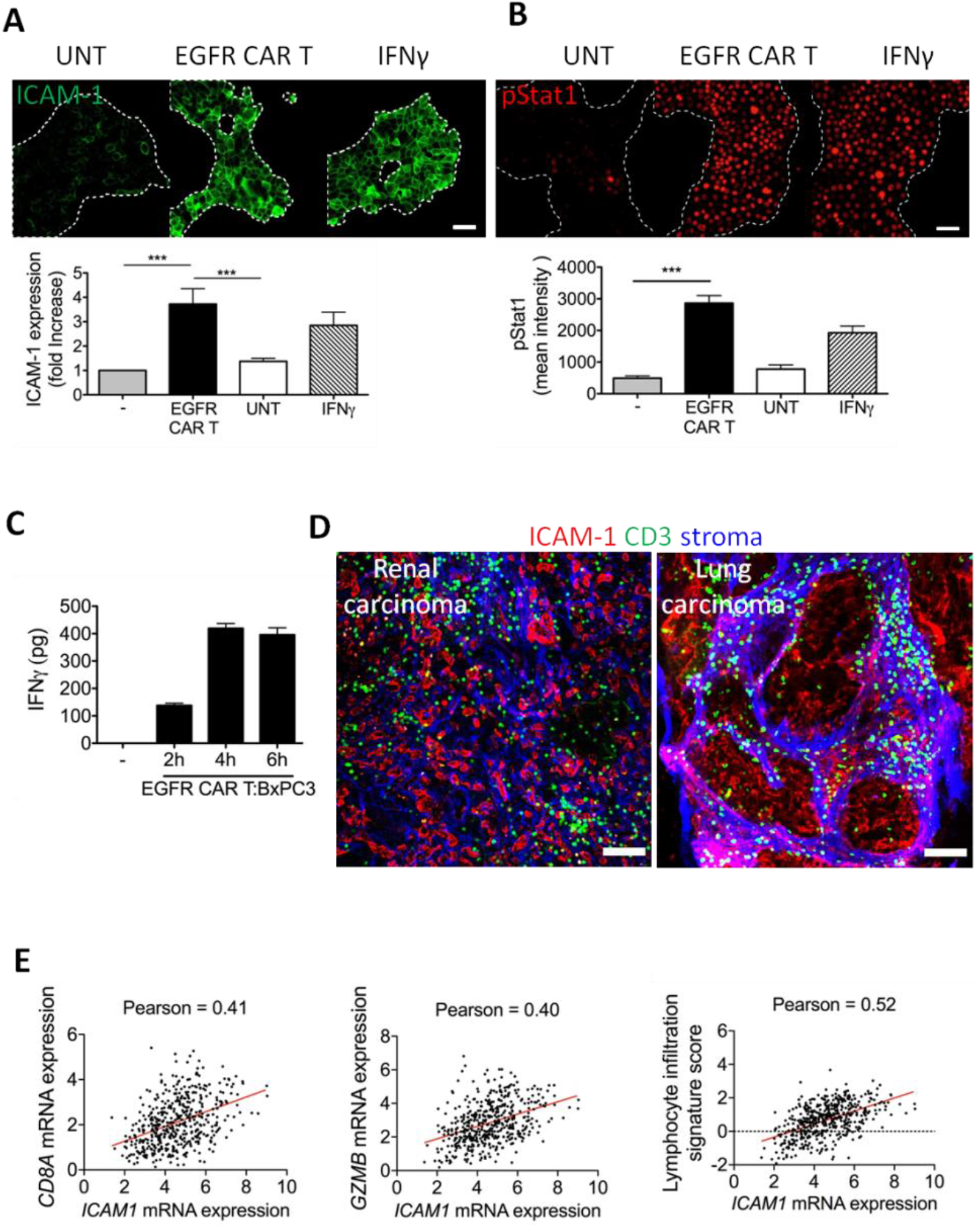
Activated EGFR CAR T cells mediate ICAM-1 expression by carcinoma cells. **(A and B, upper panels)** Immunofluorescence microscopy of BxPC3 cells left untreated, cultured with EGFR CAR T cells for 4 h at the Effector: Target ratio of 5:1 or treated with 10ng/ml IFNγ for 4 h and stained for ICAM-1 or phosphorylated STAT1 (pSTAT1). Dashed lines delineate BxPC3 cell clusters. Scale bar: 50 μm. (**A and B, bottom panels**) Quantification of ICAM-1 and pSTAT1 fluorescent intensity. Mean ± SEM; n = 4 independent experiments; Student test: ***P < 0.001. (**C**) Amount of IFNγ in the supernatant of EGFR CAR T - BxPC3 co-cultures determined by ELISA. Mean ± SEM; n = 3. (**D**) Immunofluorescence microscopy of vibratome sections from human renal and lung carcinomas stained for ICAM-1, CD3 and fibronectin. Scale bar: 100 μm (**E**) Pearson correlation coefficients of ICAM-1 mRNA expression from the TCGA lung squamous cell carcinoma dataset with CD8A (left), GZMB (middle) and a lymphocyte infiltration signature (right). R values are shown in the plots. P*** < 0.001 for all plots.

Overall, these data suggest that activated T cells are able to induce carcinoma cells to upregulate ICAM-1.

### CAR T cell-induced ICAM-1 expression on tumor cells facilitates lymphocyte entry into tumor islets

We next investigated the consequences of CAR T cell-induced ICAM-1 expression on carcinoma cells. We hypothesized that this process creates a positive feedback loop by promoting CAR T cell activation and migration. To address its kinetics, we followed the distribution of EGFR CAR T cells at different times (30 min, 2 h and 4 h) after adding them onto BxPC3 tumor cells (**Figure 5A**). Our data show that at 30 min and 2 h, most of the EGFR CAR T cells were located at the periphery of the tumor cell clusters, confirming our previous finding illustrated in Figure 2. However, 4 h after plating, EGFR CAR T cells were preferentially found within tumor cell clusters. This shows that CAR T cells progressively acquire the ability to infiltrate deeper tumor cell regions. A similar infiltration into tumor islets was observed with CD20 CAR T cells and CD20-expressing BxPC3 (**Figure S5A**). The recruitment of CAR T cells is even more obvious when selecting the most adherent cells by a light washing before taking the images. At 4 h post-plating, few untransduced T cells adhered to tumor cells within the islets. In contrast, numerous EGFR CAR T cells were found in tumor cell clusters (**Figure 5B**). To better mimic a carcinoma organization, BxPC3 tumor cells were co-cultured with human immortalized hepatic stellate cells. In this model recreating a structure of tumor cell regions surrounded by mesenchymal cells, EGFR CAR T cells interact first with peripheral tumor cells before infiltrating the center of tumor islets (**Figure S5B**). This cell recruitment was again associated with an ICAM-1 upregulation by BxPC3 tumor cells.

**Figure 5:**
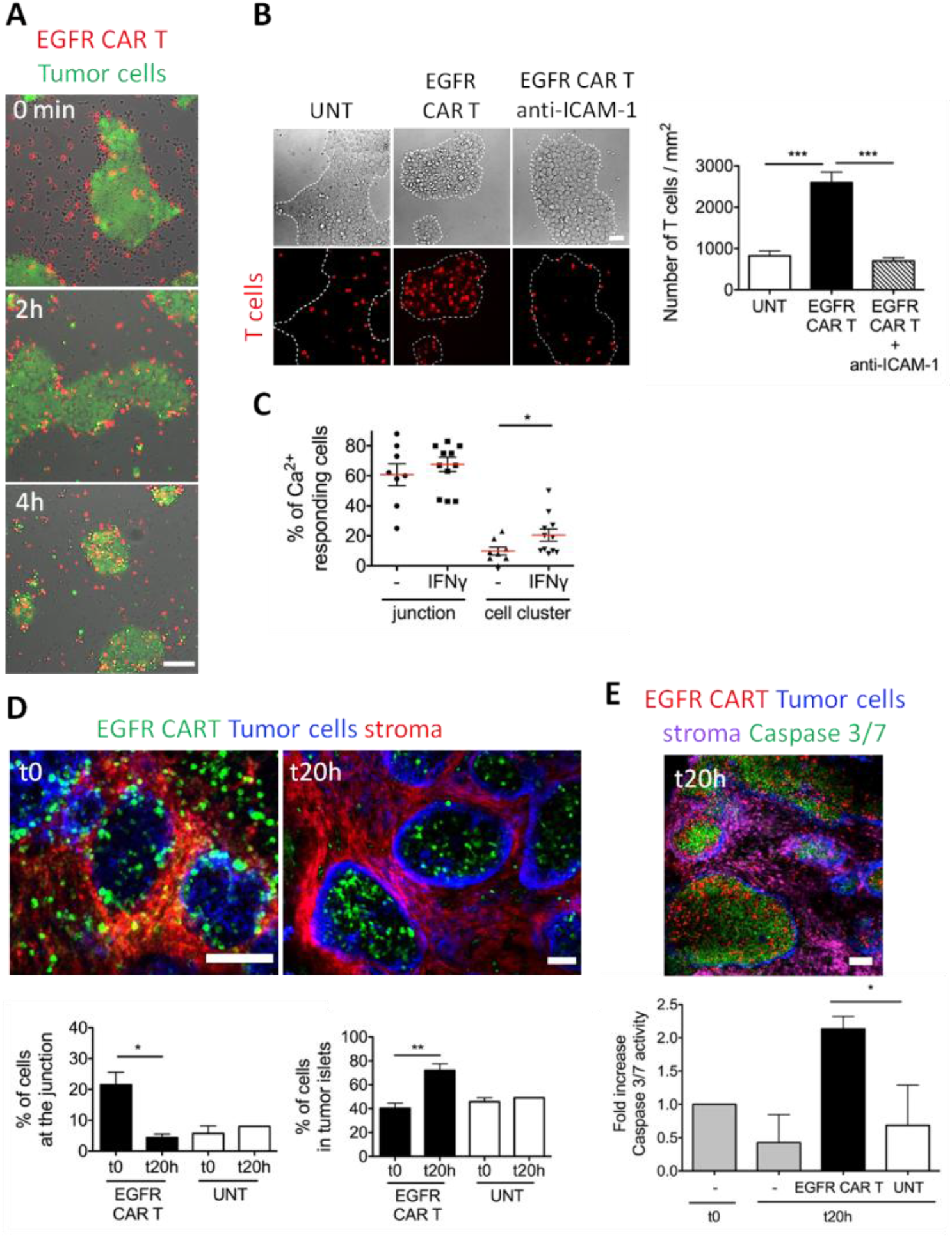
CAR T cell-induced ICAM-1 expression by tumor cells facilitates lymphocyte entry into tumor islets. (**A**) Snapshots showing EGFR CAR T cells in relation to BxPC3 tumor cells at 0, 2 and 4 h. Scale bar: 50 μm. (**B**) Untransduced and EGFR CAR T cells were allowed to interact for 4 h with BxPC3 tumor cells pretreated (right panel) or not with 10 μg/ml monoclonal anti-ICAM-1 Ab (clone HA58). Pictures were then captured after washing. Scale bar: 50 μm. (**B, right**) Concentration of EGFR CAR and untransduced T cells in BxPC3 tumor cell regions. BxPC3 cells were treated or not with anti-ICAM-1 Abs. (**C**) Proportion of Ca^2+^ responding fura-2-loaded EGFR CAR T cells at the junction and in clusters of BxPC3 cells pretreated or not with 10 ng/ml of IFNg for 4 h. (**D, upper**) Confocal images of EGFR CAR T cells distribution in slices from BxPC3 tumors stained for EpCAM (tumor cells) and fibronectin (stroma) at 0 and 20 h. (**D, lower**) Proportion of EGFR CAR and untransduced T cells at the tumor-stroma junctions (left) and in tumor islets (right) of BxPC3 tumors at 0 and 4 h, represented as mean ± SEM; n = 3 independent experiments; Student test: *P < 0.05 and **P < 0.01. (**E, upper**) A confocal image of a BxPC3 tumor slice 20 h after culture with EGFR CAR T cells in the presence of a green-fluorescent caspase 3/7 probe. Slices were stained for EpCAM (tumor cells) and fibronectin (stroma) before taking images. Note the appearance of apoptotic (green) cells in tumor cell regions enriched for CAR T cells. (**E, Lower**) Quantification of cell killing in BxPC3 tumor slices, left untreated or cultured or not with untransduced and EGFR CAR T cells for 0 and 20 h. Error bars represent SEM of 3 biological replicates. Student test: *P < 0.05. Scale bars of panels D and E: 100 μm.

Next, we performed experiments with blocking anti-ICAM-1 antibodies, and investigated the consequences on the recruitment of CAR T cells in BxPC3 tumor islets. As predicted, the specific blocking of ICAM-1/LFA-1 interactions prevented EGFR CAR T cells infiltration within tumor cells regions (**Figure 5B**). This strengthens the idea that ICAM-1 expression induced in tumor cells by activated T cells promotes their recruitment. A CAR T cell entry inhibition could also be observed when IFNγ was neutralized with an anti-IFNγ antibody (data not shown).

ICAM-1 not only promotes CAR T cell entry into tumor islets but also renders central BxPC3 and EG-I1 carcinoma cells more prone to activate CAR T cells, as evidenced by an increased percentage of Ca^2+^ responding CAR T cells on IFNγ-treated tumor cells (**Figure 5C and S5C**). Moreover, EGFR CAR T cells plated onto slices from ICAM-1-positive human renal tumors also exhibited Ca^2+^ responses, unlike untransduced T lymphocytes (**Figure S5D and Movie S9**).

We confirmed this two-step entry process - initial CAR T cell peripheral activation followed by an entry - on BxPC3 tumors. In line with our previous results, EGFR CAR T cells were preferentially enriched at the tumor-stroma junction rapidly after adding EGFR CAR T cells to BxPC3 tumor slices (**Figure 5D**). 20 h later, the situation was drastically different, since most CAR T cells were found within tumor islets. In addition, we found that EGFR CAR T cells at 24 h exhibited a potent ability to kill tumor cells as evidenced by a pronounced caspase 3/7 activity monitored in BxPC3 tumor slices using a fluorescent dye (**Figure 5E**).

Altogether, as summarized in **Figure 6**, these experiments suggest that the initial CAR T cell activation at the outer edge of the islets triggers the expression of ICAM-1 on tumor cells in an IFNγ-dependent process. ICAM-1 then promotes the progressive entry of CAR T cells into tumor islets that can then exert their cytotoxic activity against tumor cells.

**Figure 6:**
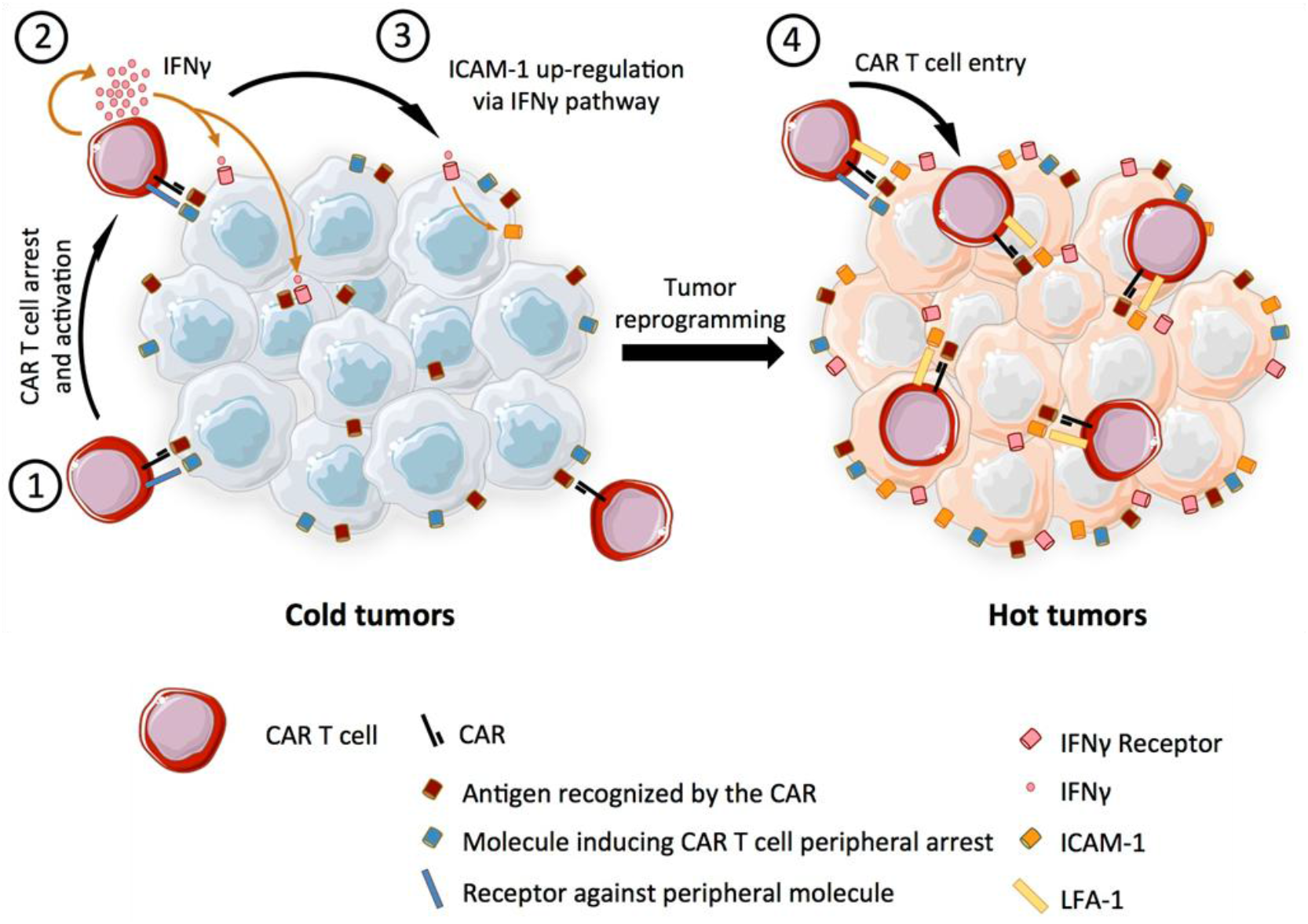
Proposed model of CAR T cell entry into tumor islets. A two-step process enables CAR T cells to infiltrate tumor islets. At first, CAR T cells interact with carcinoma cells located at the periphery of tumor islets (1). Activated CAR T cells subsequently produce IFNγ (2) which upregulates ICAM-1 by tumor cells (3). ICAM-1 expression by tumor cells facilitates CAR T cell infiltration into tumor islets.

## DISCUSSION

In this study, advanced imaging techniques applied to different tumor models including fresh human tumor slices allowed us to unravel major determinants that regulate the migration and activation of CAR T within tumor islets.

Imaging approaches have recently been used to explore the kinetics of CAR T cell activation and immune synapse formed with targets cells (12–16). However, most analyses were performed with individual tumor cells, therefore in the absence of the architecture found in human carcinomas with compact tumor islets surrounded by a stroma. Herein, using tumor models mimicking more closely carcinoma structures, we demonstrate that EGFR CAR T cell trafficking into tumor islets is regulated by a two-step process summarized in **Figure 6**. An initial CAR T cell activation at the tumor cell periphery is followed by a progressive entry into the center. The first step is an increase in ICAM-1 expression by tumor cells induced in an IFNγ-dependent pathway by an initial CAR T cell activation at the periphery of the islet. The second step is the progressive infiltration of CAR T cells within tumor islets.

Our results, obtained in two *in vitro* carcinoma cell line models and confirmed in *ex vivo* tumor slices from human lung and renal carcinomas, indicate that the spatial organization of carcinoma cells is important and dictates the responsiveness of EGFR CAR T cells. Only tumor cells located at the periphery of tumor islets are initially permissive to CAR T cell interaction and activation. Using an ultra-high content imaging approach, we found that unlike EGFR which was homogenously distributed, several proteins including the β4 integrin CD104 were enriched at the outer edge of tumor islets. Neoplastic cells within individual carcinomas often exhibit considerable phenotypic heterogeneity in their epithelial versus mesenchymal-like cell states. Interestingly, a recent study indicates that after an epithelial-mesenchymal transition, tumor cells are particularly sensitive to NK cell killing, again suggesting that tumor cells are not equal in term of immunosurveillance (17). Based on our results, we propose that these cancer cell subsets with mesenchymal features, located at the periphery of tumor islets are more prone to stimulate CAR T cells through the expression of one or several proteins that remain to be identified

In addition to specific properties of tumor cells located at the islet-stroma border, one should consider the possible role of adhesive structures present in the stroma, which could also prime T cells. Indeed, adhesion has been shown to pre-activate T cells for better responsiveness to TCR stimulation (18). In this line, CAR T cell adhesion-primed in the stroma would form productive conjugates with carcinoma cells facing this compartment. The exploration of the molecules and mechanism by which CAR T cells are initially activated at the periphery of tumor islets will be an important topic to address in the future.

Our study highlights the critical role of the adhesion molecule ICAM-1. We have shown in several models that ICAM-1 expression on malignant cells - both leukemias and carcinomas – regulates the interaction of CAR T cells with their targets as well as their migration into tumor islets. Remarkably, analyzing malignant B cells from CLL patients we found that the percentage of activated CAR T cells was positively correlated with the amount of ICAM-1 on cancer cells (figure 3B). Moreover, EGFR CAR T cells were unable to get activated and to arrest onto carcinoma cells located at the center of the islets that weakly expressed ICAM-1. Such tumor cells had a strong EGFR expression. Thus, the expression of the target antigen on carcinoma cells is not sufficient to activate CAR T cells. Adhesion molecules such as ICAM-1 play a critical role. Our results are in line with previous studies demonstrating the importance of ICAM-1 in the formation of a productive synapse between T cells and either antigen-presenting cells or target cells or between CD19 CAR T cells and malignant B cells from acute lymphoblastic leukemia patients (19–22).

Down-regulation of adhesion molecules, including ICAM-1, on tumor-associated endothelial cells is an effective mechanism by which tumors may prevent immune cell trafficking into the tumor site (23–25). Based on our data and others, we propose that the absence of ICAM-1 on tumor cells, especially in non-inflamed tumors, is an element that has to be taken into account in the therapeutic failure of CAR T cells observed in a number of malignancies. Recent studies have begun to identify reasons for such a failure including an immunosuppressive tumor environment and poor T cell survival in the tumor (26). We consider that ICAM-1 expression on tumor cells should be considered as one of the predictive markers for CAR T cell efficacy in human tumors.

Our data also indicate that when abundant enough, activated CAR T cells can overcome an initial low density of ICAM-1 by inducing its expression by tumor cells through the production of IFNγ. In other words, large enough numbers of CAR T cells can convert an immune cold into a hot tumor that becomes permissive to lymphocyte trafficking into tumor islets.

In response to the attack of tumor-specific T cells, tumor cells react by a series of modifications. Some of these have been well characterized, like the upregulation of immunosuppressive molecules, including PD-L1, in a process named ‘Adaptive-immune resistance’ of tumor cells (27). We have illustrated here another aspect characterized by an increased adhesion and attraction of CAR T cells. This is also consistent with previous findings showing that chemokines such as CXCL9, 10 and 11 are also upregulated when tumor cells are exposed to effector T cells (28). Our data demonstrate that a large number of CAR T cells in contact with carcinomas led to ICAM-1 upregulation by tumor cells. In addition, we show that in human lung and renal tumors ICAM-1 transcripts are correlated with a T cell signature. Most of these modifications are dependent on inflammatory cytokines among which IFNγ plays a key role. Nevertheless, it should be noted that IFNγ can exert both antitumour and pro-tumour functions, depending on the cellular, microenvironmental and/or molecular context (29).

Our results demonstrating the importance of this inflammatory cytokine in controlling the recruitment of CAR T cells in tumor islets is reminiscent of studies performed in auto-immune contexts. For example, T cell trafficking to inflamed pancreas during diabetes is dependent on IFNγ (30).

One of the challenges in cancer immunotherapy is to design strategies targeting ‘cold’ tumors unresponsive to most treatments including immune checkpoint inhibitors. The CAR T cell approach has been envisaged for the treatment of cold tumors due to the fact that these tumors are usually immunologically ignorant with low level of MHC expression (31). Our data support this notion. However, to be effective, a large number of T cells would have to reach cancer cells as tumor cell reprogramming, at least in our *in vitro* condition, occurs when 5 times more CAR T cells are admixed with tumor cells. We have previously evidenced several obstacles within the tumor environment, namely the extracellular matrix and macrophages, within the tumor environment hindering T cells in their displacement and ability to contact tumor cells (6, 7, 32). Thus, a combination of CAR T cells with approaches targeting these cells and elements should be envisioned in ‘cold’ tumors to enable lymphocytes to reach malignant cells in large numbers, sufficient to reprogram tumor cells.

Alternatively, tumor sensitization with IFNγ should be considered. IFNγ has been used in the 1990s as a single agent cancer immunotherapy in several prospective randomized trials but with limited evidence of clinical benefit (33). More recently, interest in IFNγ has resurged as investigators have uncovered possible synergy with other treatments and anti-tumoral effects. As an example, it has recently been shown that systemic subcutaneous injection of IFNγ in cancer patients with synovial sarcoma, a type of cold tumor, increases T cell infiltration within tumors (34).

Apart from IFNγ, other approaches have been used to increase ICAM-1 expression in cells within the tumor environment. One can mention systemic thermal therapy that was associated with IL-6 production leading to ICAM-1 expression by the tumor blood vessels thus favoring the entry of T cells to the tumor site (35).

While combination therapies gain more and more importance in oncologic treatments, CAR T approaches have not yet been combined with other drugs with the intent of augmenting lymphocyte sensitivity to low ICAM-1 levels. Based on our data, any strategies able to upregulate ICAM-1 expression on cancer cells should be considered as a new therapeutic approach to improve CAR T functionality in solid tumors.

## METHODS

### Human tumors and tumor cells

Fresh lung tumors were obtained from anonymized patients diagnosed with clinical stage I–III non-small cell lung cancer and who underwent primary lobectomy or pneumonectomy. Tumors with very high level of autofluorescence of fibers precluding imaging of T cells, with an unclear distinction between tumor cell regions and stroma were discarded after visual inspection of several microscopic fields. Fresh renal tumors were obtained from anonymized patients diagnosed with clear cell renal cell carcinoma and who underwent laparotomy. Non-fixed fresh tumors obtained after resection were rapidly transported to the laboratory in ice-cold RPMI 1640. Experiments were performed with tumor specimens obtained 2–24 h after tumor resection. CLL blood samples were obtained from untreated patients after informed consent and validation by the local research ethics committee from the Avicenne Hospital (Bobigny, France), in accordance with the Declaration of Helsinki.

### Cell lines

Human cell lines Raji from the EBV-positive Burkitt’s lymphoma and BxPC3 derived from pancreatic adenocarcinoma cells were maintained in culture in complete RPMI medium, including 10% fetal bovine serum (FBS), 50 U/mL penicillin, 50 μg/mL streptomycin. The EGI-1 tumor cell line, derived from extrahepatic biliary tract and kindly provided by Dr. L. Fouassier (INSERM, Saint-Antoine research center, Paris, France) was maintained in culture in DMEM supplemented with 1 g/L glucose, 10 mmol/L HEPES, 10% FBS, 50 U/mL penicillin, 50 μg/mL streptomycin. The stromal cell line hTERT-HSC, derived from human activated hepatic stellate cells were kindly provided by Dr. L. Fouassier and cultured in DMEM supplemented with 4.5 g/L glucose, antibiotics and 10% FBS and 2% FBS, respectively.

BxPC3 cells expressing CD20 were obtained by infection with a lentivirus encoding human CD20-IRES-GFP, followed by cell sorting.

### Xenograft models

Human B Raji lymphoma and human pancreatic BxPC3 tumors were obtained by transplantation of 10.10^6^ cells subcutaneously in the flank of immunodeficient NSG mice (Charles River). After 3 weeks, mice were sacrificed, tumors removed and processed following the same protocol described for human biopsies. Mice were maintained in the Cochin Institute specific-pathogen-free animal facility. Animal care was performed by expert technicians in compliance with the Federation of European Laboratory Animal Science Associations.

### Generation of CAR T cells

Lentiviral transfervector plasmids encoding second generation anti-EGFR CARs, were generated by cloning scFv sequences derived from Nimotuzumab or Cetuximab into the CAR expression cassette encoding CD8 hinge and transmembrane domains, 4-1BB costimulatory, and CD3z signaling domains and a 2A site followed by a truncated LNGFR or mCherry for expression monitoring. Generation of transfervectors encoding the CD20 CAR T cells was described previously (36). VSVg-pseudotyped lentiviral particles were produced in HEK293T packaging cell line using a 2^nd^ generation plasmid system. PBMCs were isolated from Buffy coat by Pancoll (PAN-Biotech) density gradient centrifugation. T cells were isolated from PBMC using the Human Pan T cell isolation kit (Miltenyi Biotec) and subsequently activated by TransAct (Miltenyi Biotec). Activated cells were cultivated in TexMACS media in presence of 10 ng/ml IL-7 and IL-15, respectively and transduced on day 1 using an MOI of 5 (TU/ml of lentivirus stock was determined on SupT1 cells). Cells were maintained at 0.5-2 x10^6^ cells /ml during expansion phase and cryopreserved on day 12. Transduction efficiency was determined on day 7 and 12 (and directly before the experiment) by anti-LNGFR staining and flow cytometric analysis.

### Tumor slices

Tumor slices were prepared as previously described (5, 6). Briefly, samples were embedded in 5% low-gelling-temperature agarose (type VII-A, Sigma-Aldrich) prepared in PBS. Tumors were cut with a vibratome (VT 1000S, Leica) in a bath of ice-cold PBS. The thickness of the slices was 400μm for solid tumors, and 800μm for Raji tumors. Slices were transferred to 0.4-μm organotypic culture inserts (Millicell, Millipore) in 35-mm Petri dishes containing 1 ml RPMI 1640 without Phenol Red.

### Antibodies

Antibodies for immunofluorescence included BV421-anti-human EpCAM (clone 9C4, BioLegend), AF408 or AF647-anti-human Fibronectin (clone HFN7.1, Novus Bio), AF647-anti-human CD3 (clone SK7, BioLegend), PE/Dazzle or BV421-anti-human EGFR (clone AY13, BioLegend), AF647 or PE -anti-human CD54 (ICAM-1) (REA266, Miltenyi), AF647-anti-mouse/human pY701-Stat1 (clone 4a, BD Biosciences), eFluor660-anti-mouse Gp38 (Podoplanin) (clone 8.1.1, 50-5381-82, eBiosciences). Flow cytometry experiments were performed using PE-anti-human CD54 (ICAM-1) (REA266, Miltenyi) and APC-anti-human CD20 (clone L27, BD Biosciences). Antibodies for blocking LFA-1 - ICAM-1 interaction were anti-ICAM1 (clone HA58, Biolegend) and anti-LFA-1 (clone TS1/18, eBioscience).

### CAR T cell imaging

Non-transduced and CAR T cells were stained at 37°C with either 1μM 5-chloromethylfluorescein diacetate (CMFDA; Invitrogen) for 5 minutes or 125nM calcein red orange for 10 minutes. For *in vitro* experiments, fluorescent T cells were added onto tumor cells cultured in ibidi μ-slides. An inverted widefield microscope (Eclipse TE2000-U; Nikon) preheated at 37°C was used for T cell imaging. In some experiments, tumor cells or CAR T cells were incubated at 37°C with 10μg/ml blocking anti-ICAM-1 and anti-LFA-1 antibodies for 20 min before the experiments.

For imaging CAR T cells in fresh tumor slices, 0.1.10^6^ or 2.10^6^ lymphocytes were plated onto sections. Imaging was performed with an upright spinning disk confocal microscope (Leica DM6000 FS, Yokogawa CSU-X1 head unit) equipped with a 37°C thermostated chamber. For dynamic imaging, tumor slices were secured with a stainless steel ring slice anchor (Warner Instruments) and perfused at a rate of 0.3 ml/min with a solution of RPMI without Phenol Red, bubbled with 95% O2 and 5% CO2. Images were acquired with a 25x water immersion objective (Leica, 25x/0.95 NA) or a 10X objective (Leica 10x/0.3 NA). For four-dimensional analysis of cell migration, stacks of 10-12 sections (z step = 5 or 15 μm for the 25x or 10x objectives respectively) were acquired every 30 s for 20 min, at depths up to 80 μm. Videos were made by compressing the *z* information into a single plane with the max intensity z projection of ImageJ software function. In some experiments, live vibratome sections were stained for 15 min at 37°C with the indicated antibodies diluted in RPMI without phenol red and used at a concentration of 10 μg/ml.

### Intracellular calcium measurement

Measurement of the intracellular Ca^2+^ concentration of T cells was performed as previously described (37) with modifications. In brief, CAR T cells were incubated for 30 min at 37°C with either 1μM fura-2 AM or fluo-4 AM (Life Technologies). T cells were then washed in HBSS and resuspended in TexMACS complete medium with 3% AB serum. For *in vitro* experiments, fura-2-loaded CAR T cells were added to the tumor cell layer cultured in ibidi μ-slides. Images were acquired every 10 seconds at 350 and 380nm. Emissions at 510 nm were used for the analysis of Ca^2+^ responses with MetaFluor software. Ca^2+^ values were represented as a ratio: fluorescence intensity at 350 nm/flurorescence intensity at 380 nm. CAR T cells were considered responsive when the amplitude of their responses reached at least twice that of the background.

For experiments in tumor slices, fluo-4-loaded CAR T cells were plated onto fresh tumor slices. Images were then acquired using the upright confocal spinning microscope described above. Fluo-4 fluorescence measurements were normalized by dividing the average fluorescence intensity (F) occurring during the course of the experiment to the average fluorescence intensity determined at the beginning of each experiments (F0). Fluorescence measurements in individual CAR T cells were performed using Imaris 7.4 (Bitplane AG).

### Cell death measurement in fresh tumor slices

Cell death within tumor slices was assessed using a green-fluorescent caspase 3/7 probe that binds DNA upon cleavage by caspase 3/7 (Nucview, Biotium). 2.10^6^ untransduced or EGFR CAR T cells were applied onto BxPC3 tumor slices that were subsequently incubated with 5μM of NucView dye. 20 h later, slices were rinsed, immunostained and imaged with an upright spinning disk confocal microscope.

### Immunostainings

Tumor cells co-cultured with CAR T cells for 1 to 4 h were fixed in 4% PFA for 10 minutes at 4°C and then washed in PBS and stained with anti-ICAM-1 (REA 266, Miltenyi) or anti-phosphorylated STAT1 pY701 (clone 4a, BD Biosciences) antibodies. For phospho-STAT1 intracellular staining, cells were permeabilized with ice-cold 90% methanol.

### MACSima

Immunostainings of a BxPC3 tumor slice (8-micron thick cryosection, fixed in acetone) in Fig. S2 were taken using a prototype of the MACSima™ Imaging Platform, a cyclic immunofluorescence device developed by Miltenyi Biotec (publication in preparation). The three displayed images of EpCAM-APC, EGFR-PE, and CD104-APC were obtained from different cycles of a much longer run (72 distinct immunolabelings from 24 3-color stain-erase cycles) on this single tissue slice. All immunostainings were obtained using recombinant, primary antibodies directly labeled with specific fluorophores (Miltenyi Biotec).

### Image Analysis

Image analysis was performed at the Cochin Imaging Facility (Institut Cochin, Paris). A 3D image analysis was performed on x, y, and z planes using Imaris 7.4 software (Bitplane AG). First, superficial planes from top of the slice to 15 μm in depth were removed to exclude T cells located near the cut surface. Cellular motility parameters were then calculated using Imaris. Tracks >10% of the total recording time were included in the analysis. When a drift in the x, y dimension was noticed, it was corrected using the “Correct 3D Drift” plug-in in ImageJ. CAR T cell concentration, motility and Ca^2+^ responses were quantified in different tumor regions, namely, stroma vs. tumor islets and islet - stroma junctions. These regions were identified by visual inspection of immunofluorescence images.

### Cytokine detection

The cytokines IFNγ and TNFα contained in supernatants derived from 1-4 h BxPC3 - CAR T cell co-cultures were quantified by ELISA using Abs from Biolegend.

### Flow cytometry

The membrane expression of CD20 and ICAM-1 on B cell from CLL patients was assessed by flow cytometry. Briefly, cells were then washed and stained with anti-CD20 and ICAM-1 antibodies for 20 min at 4 °C.

### TCGA data collection and analysis

We accessed RNA sequencing as upper quartile-normalized fragments per kilobase of transcript per million mapped reads using FireHose data repository (https://gdac.broadinstitute.org/), for each cancer of interest. Spearman correlation was used to quantify the association between ICAM-1 gene expression and T cell infiltration score, individually for each tumor type. The lymphocyte infiltration and Th1 signature gene expression signatures were previously defined (38).

### Statistics

Statistical analysis was carried out using GraphPad Prism (GraphPad software, USA). Significant differences between two series of results were assessed using the unpaired two-tailed Student’s *t* test. P < 0.05 was considered significant. Averages are expressed as mean ± SEM.

### Study approval

Human studies were carried out according to French law on biomedical research and to principles outlined in 1975 Helsinki Declaration and its modification. Institutional review board approval was obtained (CPP Ile de France II, #00001072, August 27, 2012).

Animal studies were approved by the animal experimentation ethics committee of Paris Descartes University (CEEA 34, 17-039) and by the French ministry of research (APAFiS #15076).

## AUTHOR CONTRIBUTIONS

C.K.M and E.D. designed the study; C.KM., S.B., S.H., J.M., N.B.T., A.Ki., L.V., E.Do., performed research; A.L., M.A., D.D. and M.S. provided human tumors and advices; C.KM., S.B., L.V., A.Ki. and ED analyzed data; N.B., A.T., and A.Ka. provided insights and advices; C.K.M and E.D. wrote the manuscript.

## ACKNOWLEDGEMENTS

We wish to thank Pierre Bourdoncle and Thomas Guilbert of the Cochin Imaging Facility (Institut Cochin, Paris) for advice and assistance with microscopes and help in data analysis, Fathia Ouaaz and Elisa Peranzoni for valuable discussions and critical reading of the manuscript, Marion Guérin for help in phospo-STAT1 staining and critical reading of the manuscript, Sandra Dapa for help in lentivirus production, Maximilian Wichert and Salomé Carcy for help in imaging experiments, Florence Cymbalista and Vincent Levy for providing CCL B cells and Laura Fouassier from providing human cell lines.

This study was supported by a funding from the European Union’s Horizon 2020 research and innovation programme under grant agreement No 667980.

## Abbreviations

CAR: chimeric antigen receptor
CLL: chronic lymphocytic leukemia
EGFR: epidermal growth factor
ICAM-1: intercellular adhesion molecule 1
IFNγ: interferon gamma
TNFα: tumor necrosis factor alpha

