## Supplemental Figures for "CAR T cell entry into tumor islets is a two-step process dependent on IFNγ and ICAM-1"

### Figure S1

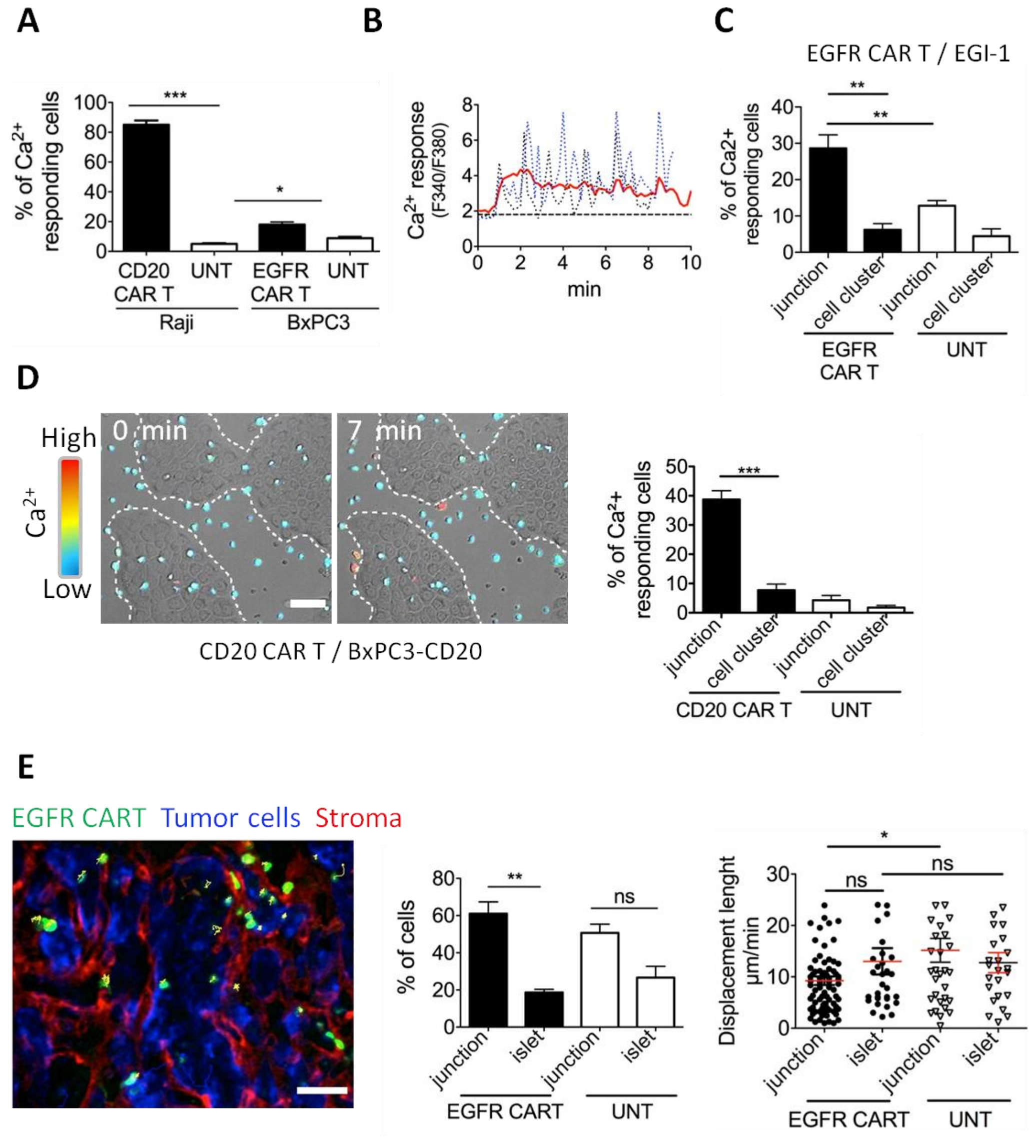


**Supplemental Figure 1. CAR T cell responsiveness is dependent on the spatial orientation of carcinoma cells**
**(A)** Percentage of $Ca^{2+}$ responding untransduced, CD20 and EGFR CAR T cells in contact with, respectively, Raji B cells or BxPC3 carcinoma cells. Results are shown as mean ± SEM; n = 100-150 cells/condition from 4-5 independent experiments; Student test: **P < 0.05, ***P < 0.001. (**B**) $Ca^{2+}$ levels of two single low affinity EGFR CAR T cells (dotted blue lines) plotted against time. Red thick lines show average $Ca^{2+}$ responses (15-20 cells.) Basal $Ca^{2+}$ levels are indicated by black dotted lines. Note the presence of $Ca^{2+}$ oscillations of large amplitude. (**C**) Proportion of $Ca^{2+}$ responding EGFR CAR T and untransduced cells at the periphery or within the tumor cell cluster of the cholangiocarcinoma cell line EGI-1. Mean ± SEM; n = 30-50 cells/condition from 3 independent experiments; Student test: **P < 0.01. (**D**) $Ca^{2+}$ response of CD20 CAR T cells contacting CD20-transduced BxPC3 tumor cells. CD20-BxPC3 cells were placed on glass coverslips the day before the experiment. CD20 CAR T cells loaded with fura 2-AM were added before image recording. **(D, left)** Snapshots of a time lapse showing $Ca^{2+}$ increases in CD20 CAR T cells after their contacts with CD20-BxPC3 tumor cells. Scale bar: 50 µm. See also **Movie S4**. (**D, right**) Proportion of $Ca^{2+}$ responding CD20 CAR T and untransduced cells at the periphery or within the tumor cell cluster of CD20-BxPC3 cells. (**E**) Concentration and migration of EGFR CAR T cells in human renal cell carcinoma tumor slices**. (E, left)** Representative images of EGFR CAR T cell distribution in a human renal cell carcinoma slice stained for EpCAM (tumor cells) and fibronectin (stroma). White dotted lines delineate tumor islets. See also **Movie S8** (**E, middle**) Proportion of EGFR CAR and untransduced T cells in the stroma, tumor-stroma junctions and tumor islets of human renal cell carcinomas, represented as mean ± SEM; n = 2 independent experiments; Student test: **P < 0.01. (**E, right**) Displacement of EGFR CAR and untransduced T cells in the stroma, tumor-stroma junctions and tumor islets of human renal cell carcinomas represented as mean ± SEM; n = 2 independent experiments; Student test: *P < 0.05. Scale bar: 50 µm

### Figure S2

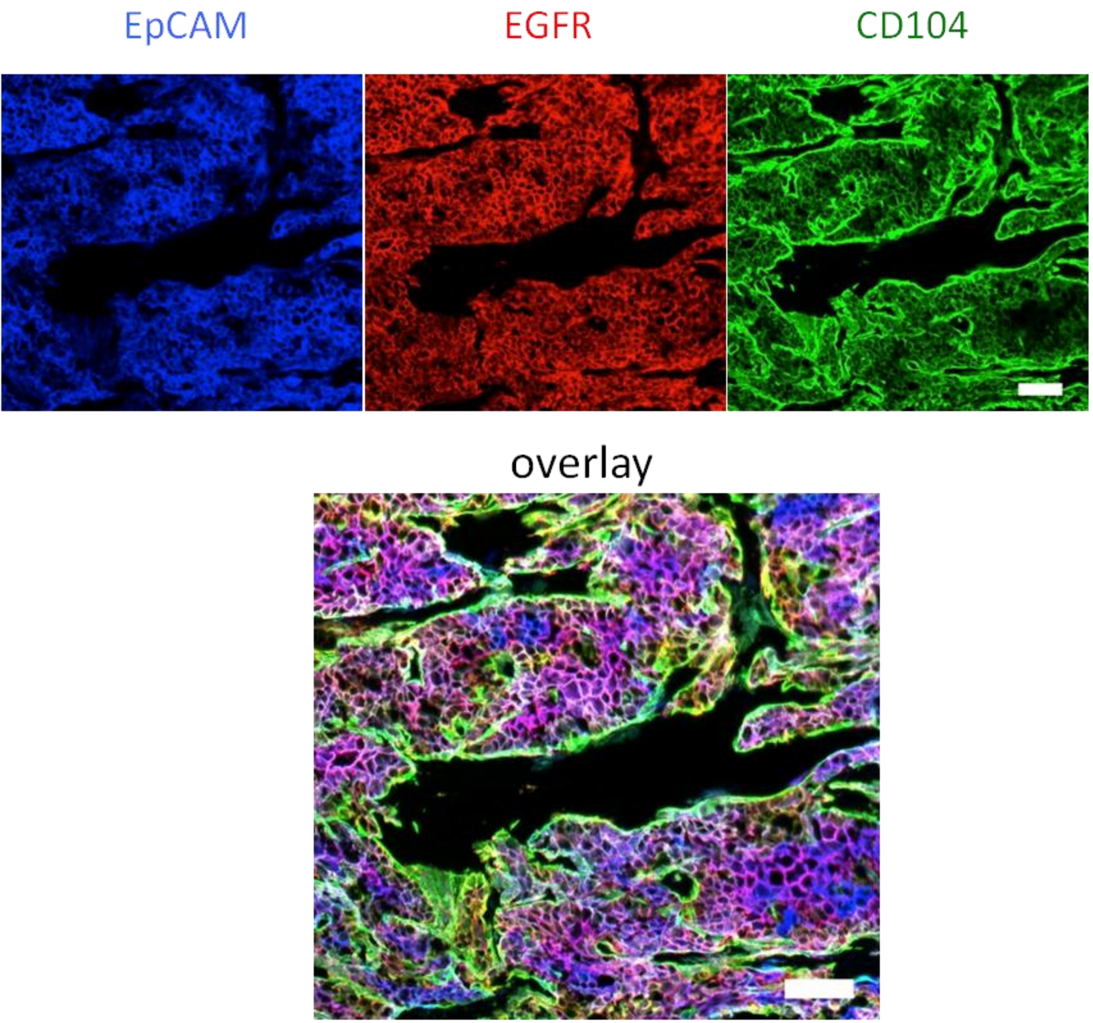


**Supplemental Figure 2. Distribution of EpCAM, EGFR and CD104 in BxPC3 tumors.**
Fixed cryosection (8-µm thick) from a BxPC3 tumor (derived from a subcutaneous tumor cell implantation into NSG mice) immunostained with the indicated antibodies. Scale bar: 100 µm.

### Figure S3

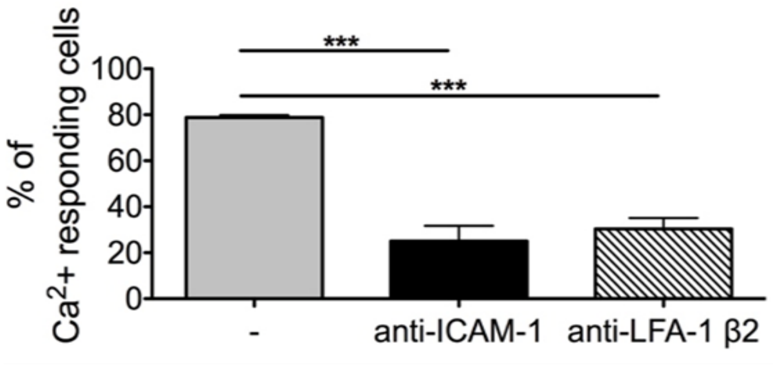


**Supplemental Figure 3. Raji B cell-induced CD20 CAR T cell $Ca^{2+}$ response is dependent on ICAM-1.**
Proportion of Fura-2-loaded CD20 CAR T cells that increases their $Ca^{2+}$ during their contacts with Raji B cells. Where indicated, ICAM-1 and LFA-1 were blocked with monoclonal antibodies (clones HA58 and TS1/18).

### Figure S4

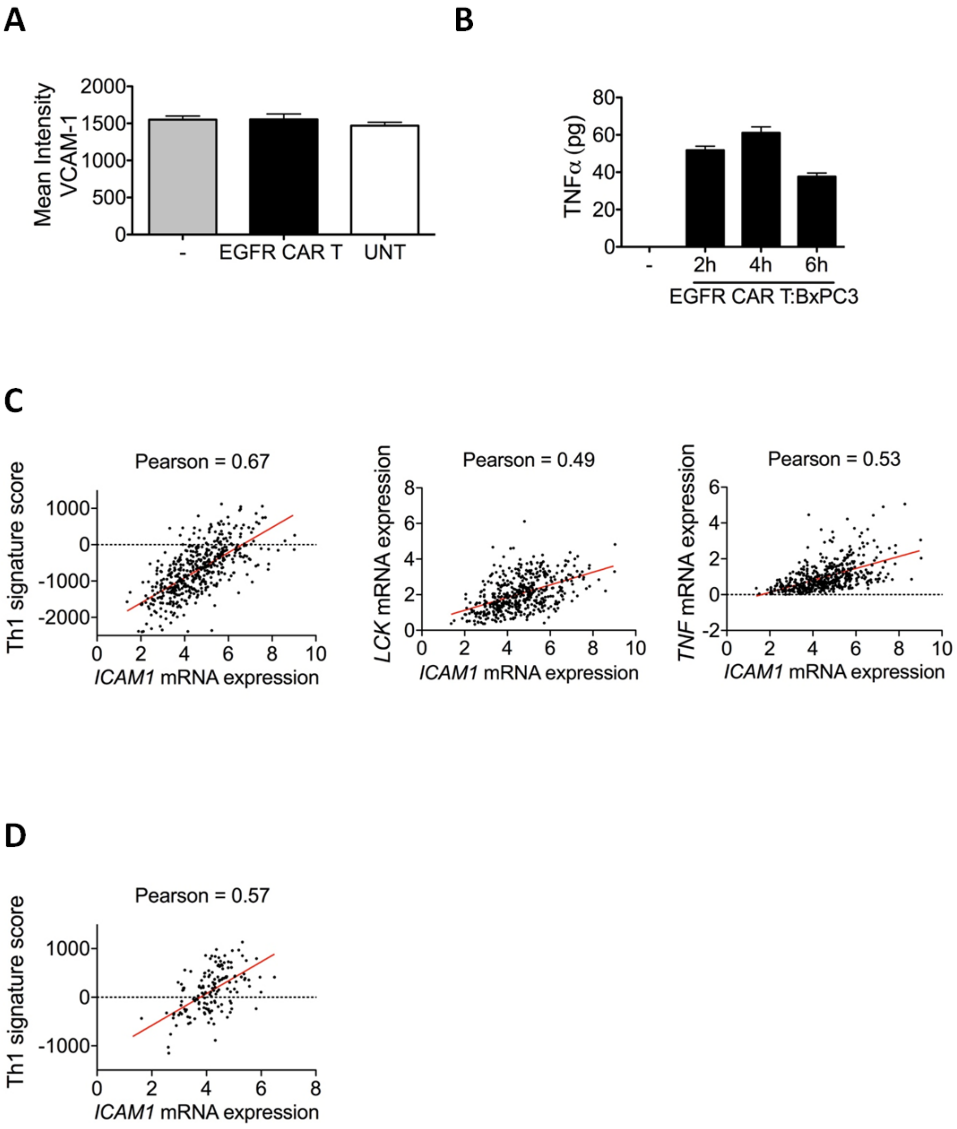


**Supplemental Figure 4. ICAM-1 expression is associated with elevated T cell infiltration in human tumors**

(**A**) EGFR CAR T cells did not mediate VCAM-1 expression by carcinoma cells. BxPC3 cells were left untreated, cultured with untransduced or EGFR CAR T cells for 4 h at the Effector: Target ratio of 5:1 and stained for VCAM-1. The histogram shows BxPC3 level of VCAM-1 fluorescent intensity. Mean ± SEM; n = 3 independent experiments. (**B**) Amount of TNFα in the supernatant of EGFR CAR T - BxPC3 co-cultures determined by ELISA. Mean ± SEM; n = 3. (**C**) Pearson correlation coefficients of ICAM-1 mRNA expression from the TCGA lung squamous cell carcinoma dataset with a Th1 signature score (left), Lck (middle) and TNF (right). (**D**) Pearson correlation coefficients of ICAM-1 mRNA expression from the TCGA pancreatic adenocarcinoma dataset with a Th1 signature score. R values are shown in the plots. P*** < 0.001 for all plots.

### Figure S5

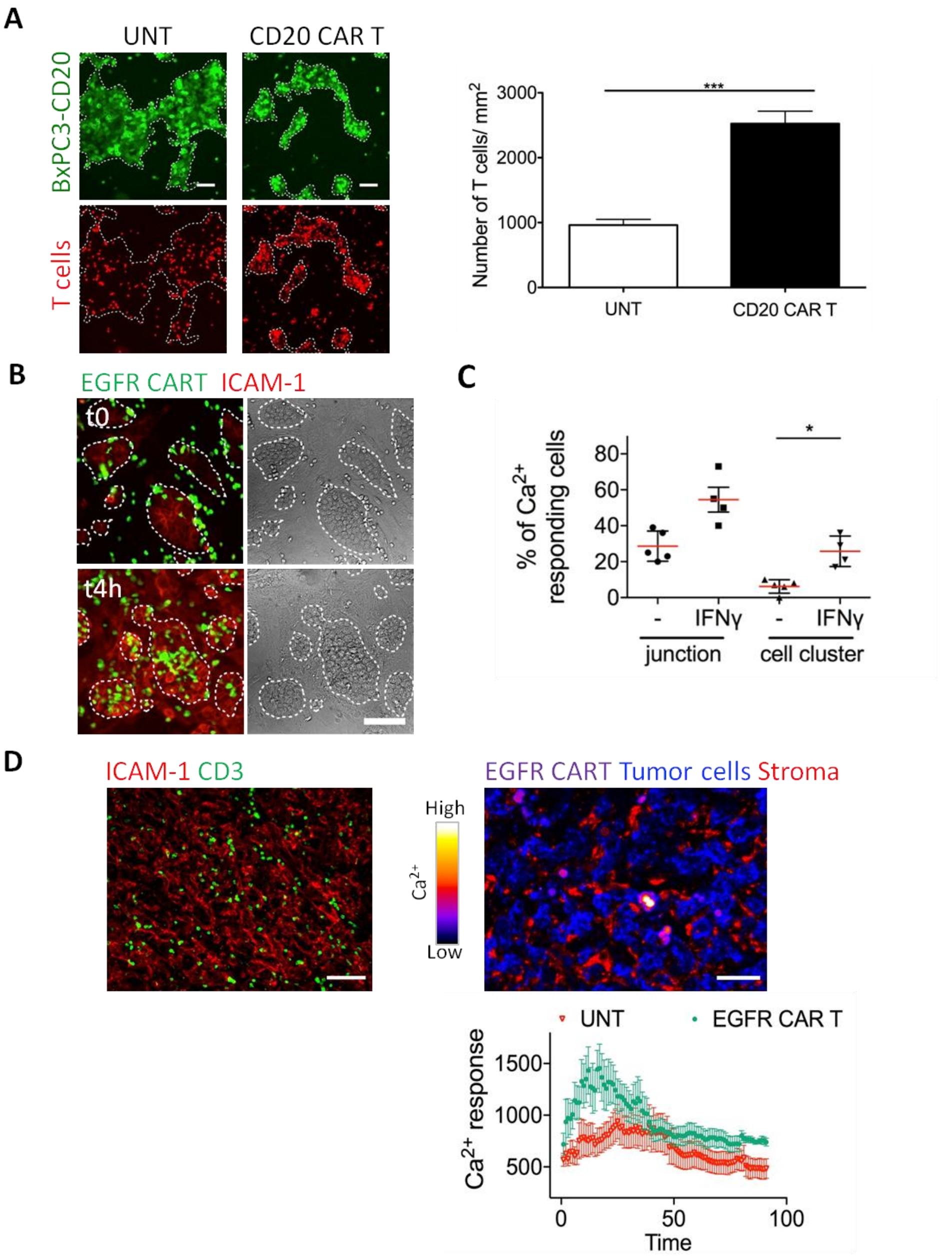


**Supplemental Figure 5. CAR T cell-induced ICAM-1 expression by tumor cells facilitates lymphocyte activation and entry into tumor islets**

(**A, left**) Snapshots showing untransduced and CD20 CAR T cells in relation to BxPC3-CD20 tumor cells at 4 h. Scale bar: 100 µm. (**A, right**) Concentration of untransduced and CD20 CAR T cells in BxPC3-CD20 tumor cell regions. Mean ± SEM; n = 3; Student test: ***P < 0.001. (**B**) Snapshots showing EGFR CAR T cells 4 h after adding them to BxPC3 - fibroblast coculture. Cells were stained for ICAM-1 before taking pictures. (**C**) Proportion of Ca$^{2+}$ responding Fura-2-loaded EGFR CAR T cells at the junction and in clusters of EGI-1 cholangiocarcinoma cells pretreated or not with 10 ng/ml of IFNg for 4 h. (**D**) Ca$^{2+}$ responses of EGFR CAR T cells in human renal tumor slices positive for ICAM-1. (**Left**) A confocal picture of a human renal tumor slice stained with the indicated antibodies. (**Right**) A confocal picture of a slice from the same tumor showing the Ca$^{2+}$ level of EGFR CAR T cells. Anti-EpCAM-1 and fibronectin Abs were used to visualize tumor cells and the stroma. See also **Movie S9**. (**Lower**) Average Ca$^{2+}$ responses of untransduced and EGFR CAR T cells (15-20 cells) in human renal tumor slices plotted against time, represented as mean ± SEM.

**Supplementary Videos**

**Movie S1. $Ca^{2+}$ responses of Fura-2-loaded CD20 CAR T cells during their interaction with Raji B cells**. Cells were imaged with a wide-field microscope. By comparison, the behaviour of untransduced T cells is shown on the right. $[Ca^{2+}]_i$ of fura-2-loaded CD20 CAR T cells is displayed in color ranging from blue (low level) to red (high level). Frame interval is 10 s. A still image is shown in Fig. 1A.

**Movie S2. $Ca^{2+}$ responses of Fluo-4-loaded CD20 CAR T cells introduced into a slice made from a Raji tumor**. Cells were imaged with a confocal spinning disk microscope. By comparison, the behaviour of untransduced T cells is shown on the right. $[Ca^{2+}]_i$ of CD20 CAR T cells is displayed in color ranging from violet (low level) to high (high level). T cell trajectories are shown white dotted lines. Frame interval is 15 s. A still image is shown in Fig. 1C.

**Movie S3. $Ca^{2+}$ responses of Fura-2-loaded EGFR CAR T cells during their interaction with BxPC3 tumor cells**. Note that CAR T cells are activated at the periphery of tumor cell regions. Cells were imaged with a wide-field microscope. $[Ca^{2+}]_i$ of fura-2-loaded CD20 CAR T cells is displayed in color ranging from blue (low level) to red (high level). Frame interval is 10 s. A still image is shown in Fig. 2A.

**Movie S4. $Ca^{2+}$ responses of Fura-2-loaded CD20 CAR T cells during their interaction with CD20-expressing BxPC3 tumor cells**. Same as in Movie S3. A still image is shown in Fig. S2D.

**Movie S5. EGFR CAR T cells accumulate and stop at the periphery of BxPC3 tumor islets**. CAR T cells were introduced into a slice made from a BxPC3 tumor that was subsequently stained for EpCAM (tumor cells) and fibronectin (stroma). Cells were imaged with a confocal spinning disk microscope. The animation represents a three-dimensional (3D) reconstruction of a sequential *z* series. Frame interval is 20 s. A still image is shown in Fig. 2B.

**Movie S6. $Ca^{2+}$ responses of Fluo-4-loaded EGFR CAR T cells in a vibratome section of a BxPC3 tumor**. The slice was stained for EpCAM (tumor cells) and fibronectin (stroma) before imaging with a confocal spinning disk microscope. $[Ca^{2+}]_i$ of CD20 CAR T cells is displayed in color ranging from violet (low level) to high (high level). White arrows indicate two CAR T cells that increase their $Ca^{2+}$ at the periphery of tumor islets. Frame interval is 15 s. A still image is shown in Fig. 1C.

**Movie S7. Distribution and migration and EGFR CAR T cells in a vibratome section of a human lung tumor**. The slice was stained for EpCAM (tumor cells) and fibronectin (stroma) before imaging with a confocal spinning disk microscope. The animation represents a three-dimensional (3D) reconstruction of a sequential *z* series. Frame interval is 20 s. A still image is shown in Fig. 2D.

**Movie S8. Distribution and migration and EGFR CAR T cells in a vibratome section of a human renal tumor**. Same as in Movie S7. A still image is shown in Fig. S1E.

**Movie S9. $Ca^{2+}$ responses of Fluo-4-loaded EGFR CAR T cells in a vibratome section of a a human lung tumor**. Same as in Movie S6. A still image is shown in Fig. S5D.
